# Seasonal and geographic variation in insecticide resistance in *Aedes aegypti* in southern Ecuador

**DOI:** 10.1101/441360

**Authors:** Sadie J. Ryan, Stephanie J. Mundis, Alex Aguirre, Catherine A. Lippi, Efraín Beltrán, Froilán Heras, Valeria Sanchez, Mercy J. Borbor-Cordova, Rachel Sippy, Anna M. Stewart-Ibarra, Marco Neira

## Abstract

Insecticide resistance (IR) can undermine efforts to control vectors of public health importance. *Aedes aegypti* is the main vector of resurging diseases in the Americas such as yellow fever and dengue, and recently emerging chikungunya and Zika fever, which have caused unprecedented epidemics in the region. Vector control remains the primary intervention to prevent outbreaks of Aedes-transmitted diseases. In many high-risk regions, like southern Ecuador, we have limited information on IR. In this study, *Ae. aegypti* IR was measured across four cities in southern Ecuador using phenotypic assays and genetic screening for alleles associated with pyrethroid IR. Bottle bioassays showed significant inter-seasonal variation in resistance to deltamethrin, a pyrethroid commonly used by the Ministry of Health, and alpha-cypermethrin, as well as between-city differences in deltamethrin resistance. There was also a significant difference in phenotypic response to the organophosphate, Malathion, between two cities during the second sampling season. Frequencies of the resistant V1016I genotype ranged from 0.13 to 0.68. Frequencies of the resistant F1534C genotype ranged from 0.63 to 1.0, with sampled populations in Machala and Huaquillas at fixation for the resistant genotype in all sampled seasons. In Machala and Portovelo, there were statistically significant inter-seasonal variation in genotype frequencies for V1016I. Resistance levels were highest in Machala, a city with hyperendemic dengue transmission and historically intense insecticide use. Despite evidence that resistance alleles conferred phenotypic resistance to pyrethroids, there was not a precise correspondence between these indicators. For the F1534C gene, 17.6% of homozygous mutant mosquitoes and 70.8% of heterozygotes were susceptible, while for the V1016I gene, 45.6% homozygous mutants and 55.6% of heterozygotes were susceptible. This study shows spatiotemporal variability in IR in *Ae. aegypti* populations in southern coastal Ecuador, and provides an initial examination of IR in this region, helping to guide vector control efforts for *Ae. aegypti*.

**Author Summary:** Mosquito control is the primary method of managing the spread of many diseases transmitted by the yellow fever mosquito (*Aedes aegypti*). Throughout much of Latin America the transmission of diseases like dengue fever and Zika fever pose a serious risk to public health. The rise of insecticide resistance (IR) is a major threat to established vector control programs, which may fail if commonly used insecticides are rendered ineffective. Public health authorities in southern coastal Ecuador, a high-risk region for diseases vectored by *Ae. aegypti*, previously had limited information on the status of IR in local populations of mosquitoes. Here, we present the first assessment of IR in adult *Ae. aegypti* to insecticides (deltamethrin, Malathion, and alphacypermethrin) routinely used in public health vector control in four cities along Ecuador’s southern coast. Observed patterns of IR differed between cities and seasons of mosquito sampling, suggesting that IR status may fluctuate in space and time. The highest overall resistance was detected in Machala, a city with hyperendemic dengue transmission and a long history of intense insecticide use. Monitoring for IR is an integral component of vector control services, where alternative management strategies are deployed when IR is detected.

## Introduction

In Ecuador, dengue, chikungunya, and Zika viruses are present and transmitted to people by the *Aedes aegypti* mosquito, causing a high burden of febrile illness in susceptible populations. This species is a particularly effective vector of these aboviruses because it has evolved to live in urban environments, lay its eggs in small containers of water in and around human dwellings, and feed preferentially on humans [1]. In Ecuador in 2016, there were 14,150 cases of dengue fever [2], 2,025 cases of chikungunya, [3] 3,531 cases of Zika fever [4]. For all three of these diseases, actual incidence in Ecuador is likely much higher than the reported figures indicate, because many cases are asymptomatic or mild, and access to laboratory diagnostics can be limited [5]. Furthermore, the co-circulation of these viruses can lead to more complex disease outcomes, and nonspecific febrile symptoms make clinical differentiation of the three diseases difficult [6]. To reduce the burden of disease, public health organizations rely heavily on vector control methods, particularly insecticide applications [7]. Additionally, individuals purchase products such as aerosols, repellents, and nets to reduce mosquito populations and prevent disease transmission at the household level, with low-income households in Machala, Ecuador, spending as much a 10% of their discretionary income on these items [5]. While vector control is considered to be the only tool available for the control of these arthropod-borne viruses, the extent to which these interventions produce significant reductions in disease burden has been difficult to ascertain [8,9].

There are four main categories of chemical insecticides regularly used for disease vector control: organophosphates, pyrethroids, carbamates, and organochlorines, with organophosphates and pyrethroids being the most widely used in Ecuador for *Ae. aegypti* control [10]. For all four of these insecticide classes, regular deployment in locations around the world has been associated with the development of insecticide resistance (IR) in targeted vector populations, resulting in resurgences of mosquito-borne diseases when vector control fails [11–13]. Although public health organizations recommend monitoring and managing IR [14], these practices are often resource-intensive, meaning that many areas do not have the capacity to conduct regular IR testing. Additionally, the largely unregulated application of insecticide treatments at the household level can influence local-scale IR. In an experimental study, exposure to commercially available aerosolized insecticides, applied in a simulation of typical household use, resulted in significant increases in genotypic and phenotypic IR in *Ae. aegypti* [15]. These results, along with field observations [16,17], indicate that fine-scale selection pressures can contribute to IR development. When resistance remains undetected, public health organizations may spend substantial time and resources applying insecticides that are ineffective [18] and may have negative environmental and human health impacts [19].

A better understanding of the spatial and temporal variability in IR would greatly enhance the ability of vector control groups to predict and mitigate IR in vector populations. Research has shown that IR status can differ dramatically across cities within a country [20–22], but further work needs to be conducted to understand the spatial scale of this variability, particularly across cities with differing vector control needs and strategies. Furthermore, most available reports from within-country spatial scales determine IR status at a single point in time, but IR exhibits temporal fluctuations that should be taken into consideration [23,24]. This study addresses a substantial knowledge gap regarding IR in *Ae. aegypti* by investigating differences in IR across both space (with four cities included in the study area) and time (with sampling occurring in three seasons), while considering both genetic and phenotypic lines of evidence to determine IR status. Furthermore, assays were conducted with multiple commonly-used insecticides, allowing for comparisons of effectiveness that can inform public health decision makers about local-scale vector-control.

In Ecuador, vector control is conducted by field workers of the Ministry of Health (MoH) in arbovirus endemic areas, as well as focal control in and around homes with suspected arboviral infections. Interventions to control adult mosquitoes include indoor residual spraying (IRS) with deltamethrin (pyrethroid) and ultra-low volume (ULV) fumigation with malathion (organophosphate). Interventions to control juvenile mosquitoes include application of an organophosphate larvicide (temephos/Abate) to containers with standing water, as well as community mobilization to eliminate larval habitats.

The objectives of this study were to evaluate overall IR status and to describe seasonal and inter-city variability in IR of Ecuadorian populations of *Ae. aegypti*, specifically testing for susceptibility to insecticides commonly used in mosquito control campaigns by the MoH. In recent work in El Oro province, researchers demonstrated resistance to deltamethrin in phenotypic assays and detected genetic mutations associated with resistance to pyrethroids in *Ae. aegypti* [25]. Beyond this small-scale study, however, IR monitoring is not regularly performed and research in this area has been limited, meaning that the prevalence of IR in these *Ae. aegypti* populations is largely unknown. Due to the high morbidity associated with the viruses transmitted by *Ae. aegypti*, as well as concerns regarding the cost and efficacy of vector control efforts, MoH leaders have indicated that there is a critical need for operational research on IR in Ecuador.

## Methods

### Ethics Statement

This study was conducted as part of a longitudinal cohort study examining social-ecological correlates of arboviral risk in Southern Ecuador. The study protocol was reviewed and approval by Institutional Review Boards (IRBs) at SUNY Upstate Medical University (IRBNET ID 4177710-25), the Luis Vernaza Hospital in Guayaquil, Ecuador, and the Ecuadorean Ministry of Health. Prior to the start of the study, all adult participants (18 years of age or older) engaged in a written informed consent process. Data collection for this project was also conducted with input from local vector control and public health organizations. Sampling coordinates were stored in a secured database and all resulting data were pooled to evaluate citywide trends in the analysis, meaning identifiable individual sites were not shared.

### Study sites and sampling period

Field samples of *Ae. aegypti* were collected in four cities in El Oro, a dengue endemic province in southern coastal Ecuador (Figure 1). The cities included in this study historically range from high to low dengue case burden. In Machala (3°15’09”S, 79°57’20”W; 6m elevation), a port city and important center of agribusiness, 136 unique sites were visited. In Huaquillas (3°28’33”S, 80°13’33”W; 15m elevation), a town on the border with Peru, 142 unique sites were visited. Both of these cities are located at sea level along the coast and have endemic transmission of dengue, typically seeing high numbers of annual reported cases. In Portovelo (3°42’58”S, 79°37’08”W; 645 m elevation), 52 unique sites were visited and in Zaruma (3°41’31”S, 79°36’47”W; 1,155m elevation), 42 unique sites were visited. These are both mining towns located further inland and at higher elevations, and have limited autochthonous transmission. The discrepancies in case burden between the cities translate into differential mosquito control demands at the municipal level, making our selection of study sites a diverse backdrop for investigating IR.

**Figure 1.**
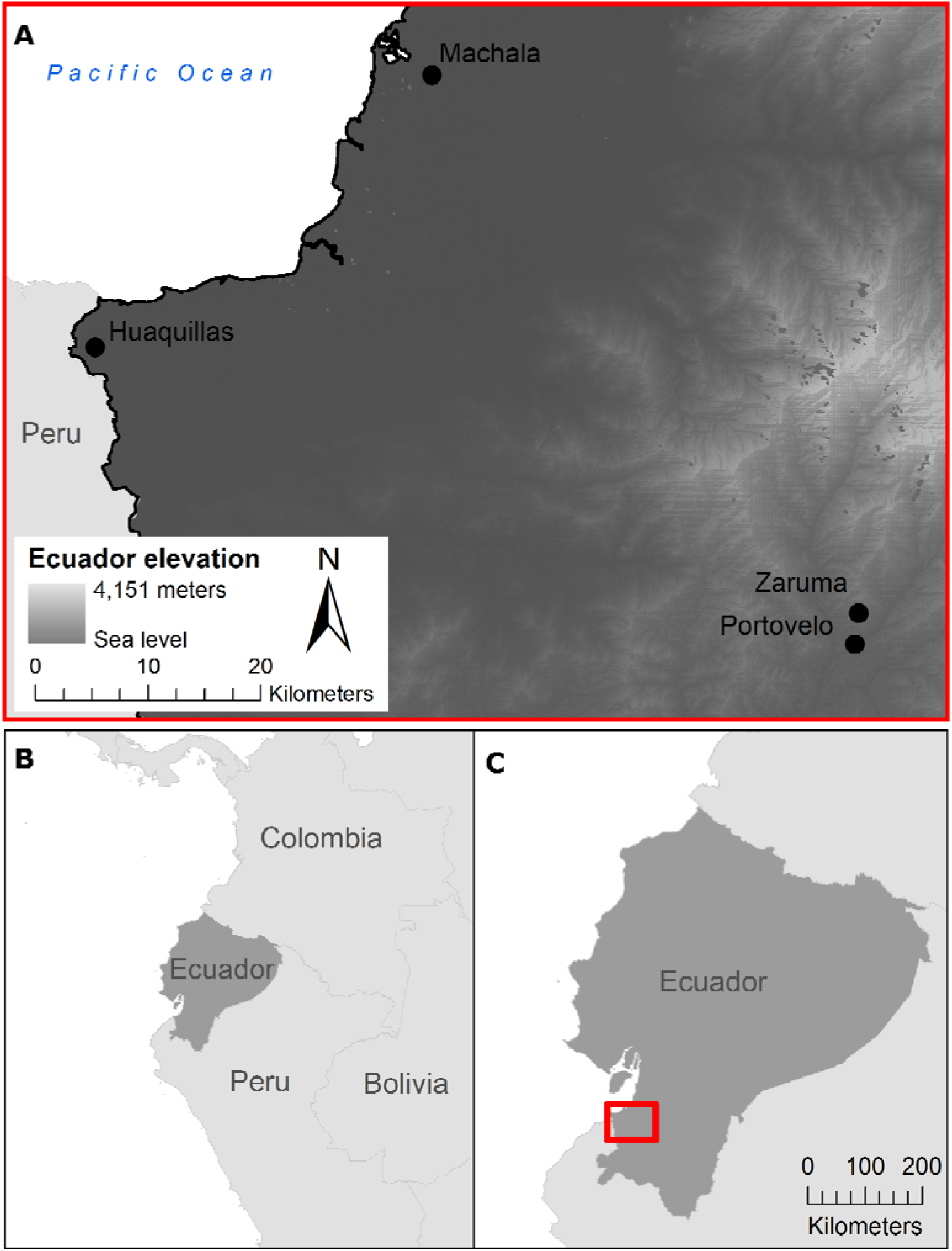
Map of the four study site cities: Machala, Huaquillas, Portovelo, and Zaruma, in southern Ecuador. Panel A shows the cities within southern Ecuador, on elevational relief; panel B illustrates Ecuador’s location along the northern Pacific coast of South America, and panel C places panel A (red bounding box) within Ecuador’s southern coastal region.

Collection of field samples was conducted in 2017 over three sampling periods: Season 1 (February 1 – April 30), Season 2 (May 1 – June 30), and Season 3 (July 1 – August 31). Seasons were explicitly chosen to collect mosquito eggs at different phases of annual arbovirus transmission in Ecuador, sampling during the peak (Season 1), decline (Season 2), and low transmission (Season 3). These collection seasons correspond with historical trends in both dengue transmission and mosquito densities in El Oro Province [26,27], and matched the observed dengue transmission pattern during the study (see S1 Figure, weekly dengue cases for each city in 2017).

### Insecticide Use Surveys

A longitudinal cohort study across the four cities included household surveys with heads of households, regarding the purchase of insecticides. Surveys were conducted from May to July 2017. We additionally collected information from the MoH in each city regarding the type, timing, and method of insecticide application over the duration of the present study.

### Egg collection and mosquito rearing

*Ae. aegypti* eggs were collected from households in the four cities using ovitraps lined with oviposition paper [28]. Two or three ovitraps were placed at each trapping site; details on the number of traps by season are shown in S2 Figure. Households were selected from ongoing surveillance study sites where the MoH designated areas of the cities as having high historic risk of dengue. These houses are distributed across each of the four cities to capture geographic variability, and are part of a larger study looking at arboviral and vector dynamics in a cohort of homes over 3 years. Ovitrapped houses for this study were purposely targeted to collect the greatest number of eggs, so ovitraps were strategically placed in areas known to have a greater abundance of eggs, representing a subset of the larger household cohort study as well as additional sites nearby. Ovitraps are a sensitive means of identifying the presence of *Ae. aegypti*, especially in areas with low vector densities [29].

Papers with eggs were collected in the field and transported to the Center for Research on Health in Latin America (CISeAL) in Quito, Ecuador, where all insect rearing and handling was performed under standard insectary conditions (28 ± 1 °C, 80% +/-10% relative humidity, 12h light; 12h darkness photoperiod). Egg hatching was achieved by placing the papers in a plastic tray containing distilled water. Larvae were fed finely ground fish flakes. Upon pupation, mosquito specimens were placed in cages for adult emergence. To increase the number of adult specimens available for experimentation, F0 adults were allowed to mate and adult female mosquitoes were blood fed on mice and allowed to oviposit. Collected eggs (F1 generation) were hatched and maintained under the aforementioned conditions. Upon pupation, F1 specimens were sorted by sex and females were used for further experimentation. Males were killed by freezing and discarded.

### Resistance monitoring

We monitored phenotypic resistance to pesticides using the bottle bioassay method as described by Brogdon and Chan [30]. Briefly, we coated glass bottles with ethanolic-insecticide solutions at established diagnostic doses: 10 μg / bottle for both deltamethrin and alpha-cypermethrin (both pyrethroids), and 50 μg / bottle malathion (an organophosphate). Each bioassay replicate consisted of four pesticide-coated bottles and one control bottle, coated only with the ethanol diluent. After the ethanol had evaporated from the bottles, groups of 15-25 female mosquitoes were introduced into each of the four treatment bottles, resulting in a total combined count of mosquitoes in the treatment bottles for each replicate ranging from 62 to 100. Mortality was recorded at 15-minute intervals. Mosquitoes were considered dead when they were incapable of flying or maintaining an upright posture on the surface of the bottle. Final mortality counts were recorded after 30 minutes of exposure.

The number of F1 females procured from field-collected specimens allowed us to perform one to three bioassay replicates for each city. The selection of pesticides (deltamethrin, alpha-cypermethrin and malathion) corresponds to pesticides typically used for vector control in Ecuador.

### Genotyping

In addition to the monitoring of phenotypic resistance, we also performed genetic screening for two mutations of the voltage-gated sodium channel gene (V1016I and F1534C) that are associated with IR to pyrethroids in *Ae. aegypti*. Both mutations have been previously described and are reported to alter the transmembrane domain of the voltage-gated sodium channel, preventing pyrethroids from binding to these sites [28,29]. Previous genotyping work has shown that *Ae. aegypti* that are homozygous for the isoleucine variant at position 1016 (I1016) are resistant to pyrethroid treatments, while those that are homozygous for the valine variant (V1016) allele are susceptible, and those that are heterozygous show intermediate resistance [33]. At position 1534, the cysteine variant (C1534) confers resistance, and the phenylalanine variant (F1534) is susceptible, while the heterozygote typically shows intermediate resistance, though there is some disagreement regarding the extent to which the heterozygous genotype confers resistance, independent of additional resistance mechanisms [32,33].

Following the aforementioned bioassays, DNA was extracted from specimens of known phenotype (susceptible/resistant to each of the pesticides used for testing). Genomic DNA was obtained using the Wizard ^®^ Genomic DNA Purification Kit (Promega Corporation, Madison, WI, USA), following the protocol established by the manufacturer. Quantification of the concentration (ng/μL) and purity (absorbance index at 260nm / 280 nm) of the obtained DNA was completed using a Nano-Drop 1000 V3.7 spectrophotometer (Thermo Fisher Scientific Inc., Wilmington, DE, USA).

Screening for the V1016I and F1534C mutations was performed as previously described [28,29], using a Bio-Rad real-time thermocycler CFX96 (Bio-Rad, Hercules, CA, USA). Results were visualized using CFX Manager Software (Version 3.1, Bio-Rad, Hercules, CA, USA). Determination of the genotype was possible through the visualization of PCR product melting curves. In the case of the V1016I mutation, a melting peak at 79°C corresponded to isoleucine (I/I – resistant mutant) and a melting peak at 85°C corresponded to a valine (V/V – susceptible wild type) (S3 Figure). In the case of the F1534C mutation, a melting peak at 85°C corresponded to cysteine (C/C, resistant mutant) and a melting peak at 80°C corresponded to a phenylalanine (F/F– susceptible wild type) (S4 Figure). Genotypic and allelic frequencies were calculated for both mutations.

Genotypic frequencies for all four cities were mapped using ArcMap© version 10.6 (Environmental Systems Research Institute, Redlands, California, USA). Inter-city and inter-seasonal differences in genotypic frequencies were tested for statistical significance (α = 0.05) using Fisher’s exact test of independence in the R software for statistical computing, version 3.4.3 (The Foundation for Statistical Computing, Vienna, Austria). When significant differences were detected, post-hoc pairwise Fisher’s tests with Bonferroni corrections were then performed to identify which cities and seasons differed significantly. The same statistical methods were used to examine phenotypic frequencies of resistance, as determined by bottle bioassays for deltamethrin, alpha-cypermethrin, and malathion, between cities for each collection season and in relation to corresponding genotypes. Additionally, populations for each city and season were assessed for Hardy-Weinberg equilibrium using the exact test in the HardyWeinberg R package [34].

## Results

### Insecticide Use

In Machala, 27.5% (14/51) surveyed homes reported purchasing pyrethroid insecticides to use at home, while 18.9% (10/53) of surveyed homes in Huaquillas reported these purchases. Of households surveyed in Portovelo and Zaruma, 45.6% (22/48) and 36.5% (19/52) respectively, reported purchase of pyrethroid insecticides for home use. Ministry of Health insecticide applications in Machala utilized a combination of deltamethrin, alpha-cypermethrin, and malathion over the duration of the study. Both Portovelo and Zaruma applied deltamethrin and malathion during the peak transmission season, followed by deltamethrin only for the following months. Huaquillas included malathion and an additional (unidentified) product during peak transmission season, followed by deltamethrin and an additional (unidentified) product in the following months (S5 Figure).

### Resistance monitoring

High levels of IR to deltamethrin, alpha-cypermethrin, and malathion were detected in all four cities, as determined by bottle bioassays. Machala had the lowest combined mortality averaged across the three seasons (18.29%, SE±5.75), indicating the highest level of resistance, followed by Portovelo (31.08%, SE±6.23) and Huaquillas (44.79%, SE±10.87). The mean mortality rate for Zaruma was 23.67% (SE±17.93) in Season 1, which was the only time period in which sufficient numbers of *Ae. aegypti* were collected for bioassays at this location.

Significant phenotypic differences in deltamethrin resistance between cities were detected in all collection seasons (Fisher’s exact test p-value<0.001) (Figure 2a). This pattern was also seen in the post-hoc pairwise tests (S1 Table). Mean mosquito mortality after 30 minutes of exposure to deltamethrin in bottle bioassays ranged from 0.70% (SE±0.6) in Machala to 18.99% (SE±0) in Huaquillas in Season 1, from 7.09% (SE±4.19) in Portovelo to 16.88% (SE±7.51) in Huaquillas in Season 2, and from 0.37% (SE±0.37) in Machala to 56% (SE±2.49) in Huaquillas in Season 3, with no mortality observed in control groups (Figure 2a).

**Figure 2.**
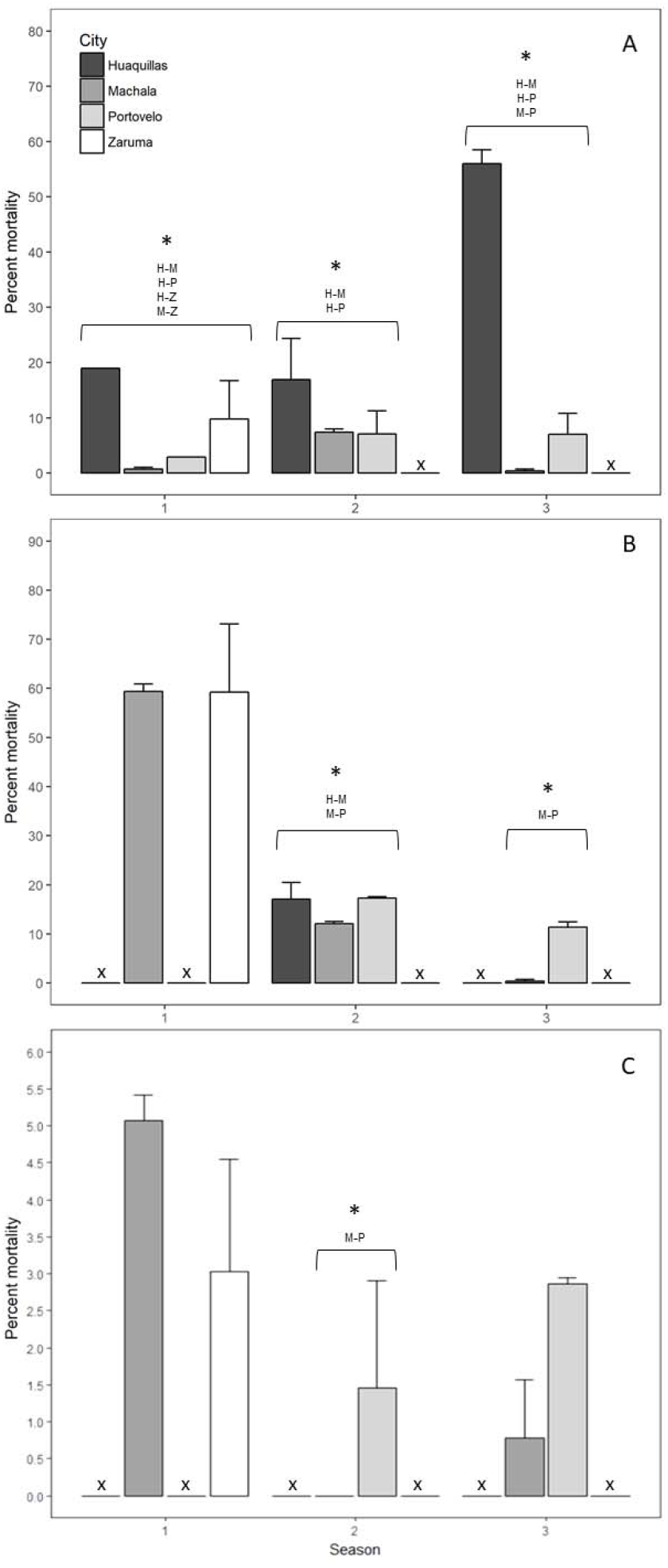
Bar plots of CDC bottle bioassay mortality for deltamethrin (A), alpha-cypermethrin (B), and malathion (C). Note different axis height for (C). Asterisks indicate Fisher’s exact test showed statistically significant differences across cities for that season. Insufficient numbers of field-collected mosquitos for bioassay within a city in a given season are indicated by ‘x’.

Mean mosquito mortality after 30 minutes of exposure to alpha-cypermethrin across bottle bioassay replicates ranged from 59.24% (SE±13.86) in Zaruma to 59.41 (SE±1.49) in Machala in Season 1, from 12.1% (SE±0.42) in Machala to 17.27 (SE±0.33) in Portovelo in Season 2, and from 0.38% (SE±0.38) in Machala to 11.39 (SE±1.03) in Portovelo in Season 3, with no mortality observed in control groups (Figure 2b). Significant differences in mean mortality rates between cities were detected in seasons two and three (Fisher’s exact test *p* value<0.001; Figure 2b), with Machala having the highest proportion of phenotypically resistant specimens in these two seasons (S2 Table). Collection counts were insufficient for alpha-cypermethrin bioassays in Portovelo for Season 1, Huaquillas for Seasons 1 and 3, and Zaruma for Seasons 2 and 3.

Mean mosquito mortality after 30 minutes of exposure to malathion in the bottle bioassays ranged from 5.07% (SE±0.35) in Machala to 3.03% (SE±1.52) in Zaruma in Season 1, from no mortality in Machala to 1.45% (SE±1.45) in Portovelo in Season 2, and from 0.78% (SE±0.78) in Machala to 2.86% (SE±0.08) in Portovelo in Season 3, with no mortality observed in control groups. Statistically significant differences in Malathion resistance were only detected in season two between Machala and Portovelo (Fisher’s exact test *p* value < 0.001). Due to low sample sizes, bioassays could not be conducted in Portovelo for Season 1, Zaruma for Seasons 2 and 3, and Huaquillas for all three seasons, and therefore further comparisons were not feasible.

Inter-seasonal variations in phenotypic resistance for each city were also assessed for statistical significance. In the deltamethrin bottle bioassays, mean mortality in Machala increased from Season 1 to Season 2, but decreased from Season 2 to Season 3 (Post-hoc Fisher’s exact test p value <0.001), while there were increases between each season in both Huaquillas (Post-hoc Fisher’s exact test p value <0.003) and Portovelo (Fisher’s exact test p value < 0.02; S3 Table). In the alpha-cypermethrin bottle bioassays, significant inter-seasonal differences were only detected in Machala, where the percent mortality decreased between each season (Post-hoc Fisher’s exact test p value < 0.05; S4 Table). Similarly, significant inter-seasonal differences in mean mortality in the malathion treatment were only detected in Machala, where mortality decreased from Season 1 to Season 2 (Post-hoc Fisher’s exact test p value < 0.05; S5 Table).

### Genotyping

Observed genotype frequencies varied significantly between cities only in Season 1 for both V1016I and F1534C alleles (Fisher’s exact test p value < 0.001; Figure 3). Pairwise post-hoc analysis revealed significant differences in the frequencies of genotypes I/I (mutant) and V/I (heterozygous) for the V1061I gene, with Huaquillas (n = 34) having a significantly higher frequency of heterozygotes compared to Machala (n = 22) and Portovelo (n = 22; Post-hoc Fisher’s exact test p values < 0.001 and 0.03, respectively; S6 Table). Portovelo and Zaruma (n = 40) differed in the frequency of I/I (mutant) and V/V (wild type) genotypes, with Zaruma having a significantly higher frequency of wild type mosquitoes during Season 1 (Post-hoc Fisher’s exact test p value < 0.05; S6 Table). Genotypic frequencies of C/C (mutant) and F/C (heterozygous) genotypes of the F1534C resistance gene differed across cities in Season 1, with Zaruma having a significantly higher proportion of the heterozygous genotype than Huaquillas and Machala (Post-hoc Fisher’s exact test p values < 0.02 and < 0.001, respectively; S7 Table). Although the frequencies of F1534C genotypes also varied significantly in the Season 3 (Fisher’s exact test p value < 0.03), conservative post-hoc analysis did not reveal any significant pairwise relationships (S7 Table).

**Figure 3.**
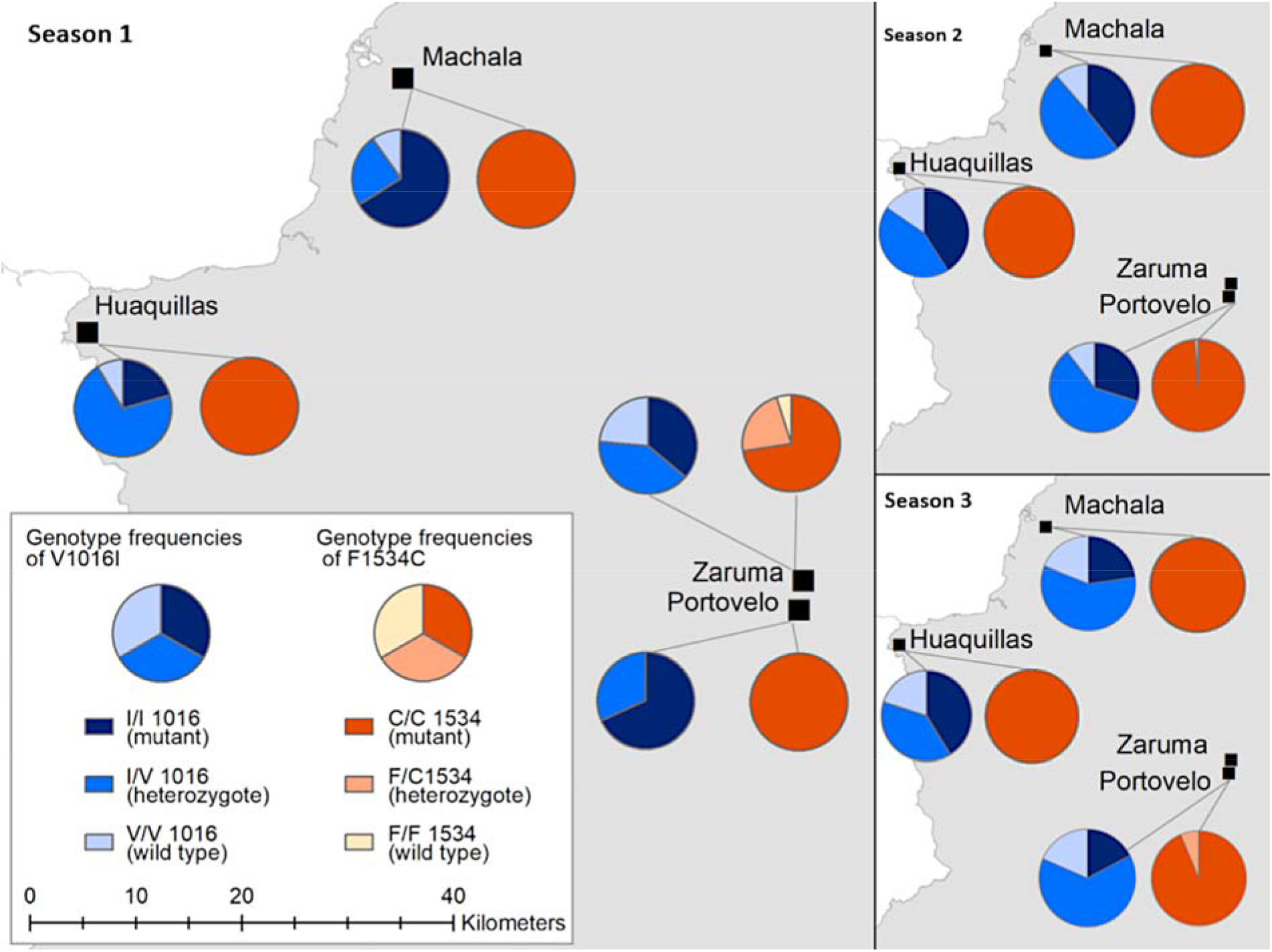
Genotypic frequencies of the V1016C and F1534C alleles at the four study site locations during Season 1 (Feburary 1-April 30, 2017).

Significant inter-seasonal variation in V1016I genotype frequencies was detected in Huaquillas, Machala, and Portovelo (Fisher’s exact test p values < 0.04, <0.001, and <0.001, respectively), though conservative post-hoc analysis did not identify significant pairwise differences in Huaquillas. In Machala, the proportion of the I/I (mutant) and V/I (heterozgote) genotypes differed significantly from Season 2 to Season 3 (Post-hoc Fisher’s exact test p value < 0.02), with the I/V genotype increasing while the I/I genotype decreased (S8 Table). In Portovelo, there were significant differences in the frequencies of the I/I and V/I genotype between Seasons 1 and Season 2 and between Season 1 and Season 3 (Post-hoc Fisher’s exact test p values < 0.05 and <0.001, respectively), with the frequency of the I/I genotype decreasing while V/I increased. Similarly, the frequency of the V/V (wild type) genotype increased significantly from Season 1 to Season 3 (Post-hoc Fisher’s exact test p value < 0.005) while the I/I genotype decreased (S9 Table). No significant inter-seasonal differences in F1534C genotype frequency were detected.

There were statistically significant associations between the resistance genotypes and phenotypic resistance results for 632 *Ae. aegypti* that were genotyped and subjected to the pyrethroid (deltamethrin or alpha-cypermethrin) bottle bioassays. Due to low sample sizes, we pooled pyrethroid assay results (Tables 1 & 2), and present the separated analyses in supplemental information (S10, S11 Tables). For mutation V1016I, while the majority of the resistant individuals (96.4%) presented the mutant or heterozygous genotype, 12 (3.6%) phenotypically resistant individuals presented the wild type (V/V) genotype that is typically associated with susceptibility to pyrethroids, and 40 (13.5%) susceptible individuals presented the genotype that typically confers resistance. Similarly, at the F1534C locus, 276 (93.2%) individuals with the mutant (C/C) genotype, typically associated with resistance, were susceptible to pyrethroid treatments. Due to low sample sizes of the wild type (F/F) for both pyrethroids, separate Fisher’s exact tests were not conducted (S6 Table). Because the two loci studied here contribute additively to resistance, the proportions of resistant and susceptible individuals for each combined V1016I and F1534C genotypes were also considered. Of the 227 mosquitoes with the mutant I/I and the mutant C/C genotype, 187 (82.4%) were resistant in the pyrethroid assay. There were twelve genotyped mosquitoes that were heterozygous at both loci, and of these, six individuals were resistant; likewise, only three genotyped specimen were homozygous wild type at both loci, and all three were susceptible (Tables 1 & 2). The V1016I genotype frequencies indicate that populations from Huaquillas and Machala during Season 1 (Fisher’s exact test p values < 0.03) and Portovelo during Season 2 (Fisher’s exact test p value < 0.02) were not in Hardy-Weinberg equilibrium (S12 Table).

**Table 1.** Occurrences of V1016I and F1534C genotypes compared to pyrethroid-resistant phenotype. Fisher’s tests indicated significant associations between genotypes and phenotypes for resistance.

| V1016I | | I/I<br>(mutant) | V/I<br>(heterozygote) | V/V<br>(wild type) | $p$-value* |
| --- | --- | --- | --- | --- | --- |
| V1016I | *Resistant* | 187 | 137 | 12 | $<0.001$ |
|  | *Susceptible* | 40 | 172 | 84 |  |
| F1534C | | C/C<br>(mutant) | F/C<br>(heterozygote) | F/F<br>(wild type) | $p$-value* |
| F1534C | *Resistant* | 328 | 7 | 1 | 0.02 |
|  | *Susceptible* | 276 | 17 | 3 |  |

\* Fisher's Exact Test

**Table 2.** Combined occurrences of V1016I and F1534C genotypes compared to pyrethroid-resistant phenotype. Fisher’s tests indicated significant associations between genotypes and phenotypes for resistance.

| V1016I/<br>F1534C | | II/CC | VI/CC | VV/CC | VV/FC | VI/FC | VI/FF | VV/FF | $p$-value* |
| --- | --- | --- | --- | --- | --- | --- | --- | --- | --- |
| V1016I/<br>F1534C | *Resistant* | 187 | 130 | 11 | 1 | 6 | 1 | 0 | $<0.001$ |
|  | *Susceptible* | 40 | 166 | 70 | 11 | 6 | 0 | 3 |  |

\* Fisher's Exact Test

## Discussion

In this study, exposure to diagnostic doses of deltamethrin, alpha-cypermethrin and malathion resulted in mortality rates below 80% after 30 minutes of insecticide exposure in all populations tested, regardless of collection season. Based on the World Health Organization’s (WHO) recommendations for assessing the significance of detected resistance [30], our results suggest that these *Ae. aegypti* populations should be considered as resistant to all the insecticides considered in our study. Furthermore, the results of the bioassays for malathion susceptibility are of particular interest, as they indicate that these populations are extremely resistant to malathion, with no population showing more than 5.07% mortality, and populations from Machala reaching values as low as 0% mortality during Season 2.

The CDC bottle bioassay used in this study is one of several methods used to detect resistance in mosquito populations. The other most commonly used method, the WHO susceptibility test, is a response-to-exposure analysis that uses insecticide-impregnated papers obtained directly from a distribution center [34]. While this pre-manufactured quality means there is greater consistency and control in the administration of the WHO susceptibility test, the CDC bottle bioassay can be conducted without specialized equipment, which often makes it the assay of choice in resource-limited settings [35]. In addition to these methods, IR can be assessed based on calculation of the lethal concentration needed to kill half of a sample of mosquitoes (LC_50_) after topical application of the insecticide of interest [36]. This method allows for the calculation of resistance ratios to quantify and compare resistance across populations; however, this test is more time-consuming and requires specialized equipment, so it is not as commonly conducted [37].

In this study, molecular characterization showed that the resistance-associated mutant alleles V1016I and F1534C are present at all the locations studied. In particular, allele F1534C was present at very high frequencies (close to 1) in all the locations studied and across all three seasons, similar to results from recent work in Mexico [35]. This suggests that this gene has been subjected to selective pressures in the past and is approaching or has reached fixation in these populations. By contrast, the results from the test for Hardy-Weinberg equilibrium for the V1016I gene and the significant inter-seasonal differences in genotype frequencies indicate that the populations are still responding to varying selective pressures and are not in a state of equilibrium. Both of these mutations typically confer resistance to dichlorodiphenyltrichloroethane (DDT) as well as pyrethroids [36,37], meaning selection for these alleles likely began with earlier widespread usage of DDT (which was used by the MoH until 1996 [38]) and has persisted as pyrethroids became more commonly used [39]. Significant distinctions in genotypic and phenotypic frequencies were not detected for many cities in this study during subsequent seasons.

It is worth noting that for some locations and seasons, periods of low vector densities in the field resulted in the collection of low numbers of mosquito eggs. Overall, collection counts were consistently low in Zaruma, a small city that has the highest elevation of the four study cities. This observation is consistent with other work that has documented a negative relationship between elevation and the probability of *Ae. aegypti* presence [40]. Similarly, collection counts during Season 2 and Season 3 were not high enough for bioassays with each of the three insecticides of interest. These seasons have historically corresponded with periods of low mosquito activity and dengue transmission in Ecuador, as observed in 2017 [26,27] This scarcity of F0 individuals translated into missing data for some cities and/or very low frequency counts in some categories, thus providing a limited basis for making quantitative comparisons.

The detection of resistance to organophosphate and pyrethroid insecticides in Ecuador is consistent with broader regional trends identified in recent years. In neighboring Peru, *Ae. aegypti* strains have been shown to be resistant to multiple organophosphates and pyrethroids in WHO susceptibility tests [41]. Similarly, pyrethroid resistance has been detected in Colombia in both CDC and WHO bioassays, though the levels of resistance vary throughout the country [21]. However, Colombian *Ae. aegypti* populations tested with both WHO and CDC bioassays were broadly susceptible to malathion, in contrast to the widespread resistance seen in the Ecuadorian *Ae. aegypti* in this study [42]. Throughout the rest of the Americas, pyrethroid resistance, as measured in studies using LC50 or percent mortality, appears to be broadly distributed, particularly in Brazil and French Guiana, though some populations in Costa Rica, Panama, and northern Colombia are still susceptible [10].

The temporal and spatial variability in the results from the bioassays highlight the importance of regularly conducting IR monitoring across multiple locations to understand the true extent of IR and make appropriate vector control decisions. For example, in Machala, the mortality rate of *Ae. aegypti* treated with alpha-cypermethrin decreased significantly from Season 1 to Season 3, indicating this population was becoming less susceptible throughout the course of the study. By contrast, the mortality rate associated with the deltamethrin assays on populations from Huaquillas increased from Season 1 to Season 3, meaning this population was becoming more susceptible over time. The impact of city-level insecticide application on IR is difficult to infer with the information we obtained from the MoH of each municipality. Portovelo did not use alpha-cypermethrin and we saw no differences in alpha-cypermethrin mortality rates across seasons. Machala used some form of deltamethrin, alpha-cypermethrin, and malathion throughout the study duration, but mortality rates differed for these insecticides by season, with mortality rates decreasing for both alpha-cypermethrin and malathion. There was some seasonal variation in method of application, strength of insecticide solution, and neighborhood coverage, which could impact the IR of our sampled mosquito populations. Continued work in this area could determine if our observed trends are due to seasonal fluctuations, differential insecticide application parameters, local-scale movements of *Ae. aegypti* populations with varying levels of resistance, or long-term, inter-annual trends. There were also statistically significant differences in mortality rates across the cities for the deltamethrin bioassays in all three seasons and in two of the three seasons for the alpha-cypermethrin bioassays. Considering this variability in the resistance phenotypes found within a single province of Ecuador, organizations involved in decision-making about insecticide applications should be cautioned against inferring the IR status of one *Ae. aegypti* population based on the status of populations in neighboring municipalities.

To better contextualize this work for appropriate vector-control decision-making, the relationships between genotypes, IR bioassay results, and actual IR status in the field should be considered. While the V1016I and F1534C mutations are known markers of pyrethroid resistance in *Ae. aegypti*, the results of this study showed that the genotypes were not perfectly predictive of resistance phenotypes, even when both the V1016I and F1534C genotypes were considered. In the recorded frequencies of genotypes versus bioassay outcomes, resistant and susceptible phenotypes were observed for each genotype, although the resistant phenotype was still statistically associated with the mutant genotypes for both genes. This is likely due to other IR mechanisms, such as metabolic detoxification processes [18] that could influence IR status in these populations; however, these mechanisms were not considered in the current study. Additionally, factors such as temperature, larval nutrition, larval density and age have been shown to influence insecticide susceptibility in *Aedes* mosquitoes, leading to discrepancies between bioassay results and the actual outcomes of insecticide treatments in the field [43]. Further work on IR in *Aedes* populations could identify and possibly reconcile differences between results from the laboratory and the field. To comprehensively evaluate IR status, future studies should also investigate the effectiveness of insecticide application methods, intensity, timing, and coverage by households and the MoH, as well as the impact of larvicides, such as temephos, which is commonly used in this study area, as well as *Bacillus thuringensis israelensis* (Bti), which is a common control method in other parts of the world.

Geographic methods, particularly spatial statistics and modelling, are well suited for understanding the patterns and drivers of IR at meso- and local scales. Employing these approaches can lead to more targeted, efficient, and sustainable vector control efforts. Future research in this area should continue to explore the spatial and temporal variability in IR among *Ae. aegypti* populations. Recent work in Yucatan, Mexico, demonstrated that IR levels could vary significantly across neighborhoods within the same city [23]. Additionally, work on the temporal dynamics of resistance could be beneficial for vector control decision-making. For example, in a study on pyrethroid-resistant, field-derived *Ae. aegypti*, researchers demonstrated that susceptibility to pyrethroids could be restored within ten generations when the selective pressure of regular insecticide treatments was removed [24]. While this experiment was conducted in a controlled environment, similar work within the context of urban environments, such as the four cities included in this study, could help calibrate timing of insecticide class rotations, allowing for better long-term management of susceptibility.

In conclusion, the *Ae. aegypti* collected in these four cities in Ecuador showed varying levels of resistance to the insecticides tested, and these measures typically changed over the course of the three seasons during which sampling took place. Regular IR monitoring should be conducted as long as insecticide applications remain an integral component of vector control activities, particularly in areas where these operations are deployed to control arbovirus transmission. Beyond this monitoring process, appropriate alternative management strategies should be deployed when IR is detected. These strategies can include biological control and community mobilization to reduce *Ae. aegypti* breeding sites.

## Acknowledgments

We are grateful to Naveed Heydari, the Ministry of Health of Ecuador, and our colleagues who facilitated data collection in the field.

## Gene ID Numbers

Gene ID number is for the NCBI Gene database: https://www.ncbi.nlm.nih.gov/gene voltage-gated sodium channel gene, *VGSC:* 5567355

## Supporting information

**S1 Table:** Post-hoc Fisher’s exact test p-values for Deltamethrin resistance between cities. Significant difference are denoted with an asterisk.

| Season | City | Huaquillas | Machala | Portovelo |
| --- | --- | --- | --- | --- |
| 1 | Machala | < 0.001* |  |  |
|  | Portovelo | 0.002* | 1.00 |  |
|  | Zaruma | 0.008* | < 0.001* | 0.75 |
| 2 | Machala | 0.006* |  |  |
|  | Portovelo | < 0.001* | 1.00 |  |
| 3 | Machala | < 0.001* |  |  |
|  | Portovelo | < 0.001* | < 0.001* |  |

**S2 Table:** Post-hoc Fisher’s exact test *p*-values for Alpha-Cypermethrin resistance between cities in the season two collection period. Significant difference are denoted with an asterisk.

| City | Huaquillas | Machala |
| --- | --- | --- |
| Machala | <0.001* |  |
| Portovelo | 0.70 | <0.001* |

**S3 Table:** Post-hoc Fisher’s exact test *p*-values for Deltamethrin resistance between seasons. Significant difference are denoted with an asterisk.

| City | Season | 1 | 2 |
| --- | --- | --- | --- |
| Machala | 2 | <0.001* |  |
|  | 3 | <0.001* | <0.001 |
| Huaquillas | 2 | 0.003* |  |
|  | 3 | <0.001* | <0.001* |
| Portovelo | 2 | <0.001* |  |
|  | 3 | <0.001* | 0.02* |

**S4 Table:** Post-hoc Fisher’s exact test *p*-values for Alpha-cypermethrin resistance between seasons. Significant difference are denoted with an asterisk.

| City | Season | 1 | 2 |
| --- | --- | --- | --- |
| Machala | 2 | <0.001* |  |
|  | 3 | <0.001* | 0.05* |
| Portovelo | 3 |  | 0.24 |

**S5 Table:**
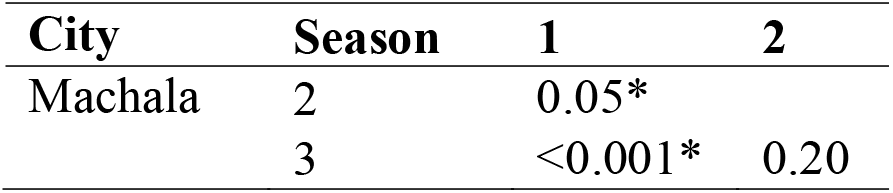
Post-hoc Fisher’s exact test *p*-values for Malathion resistance between seasons. Significant difference are denoted with an asterisk.

**S6 Table:**
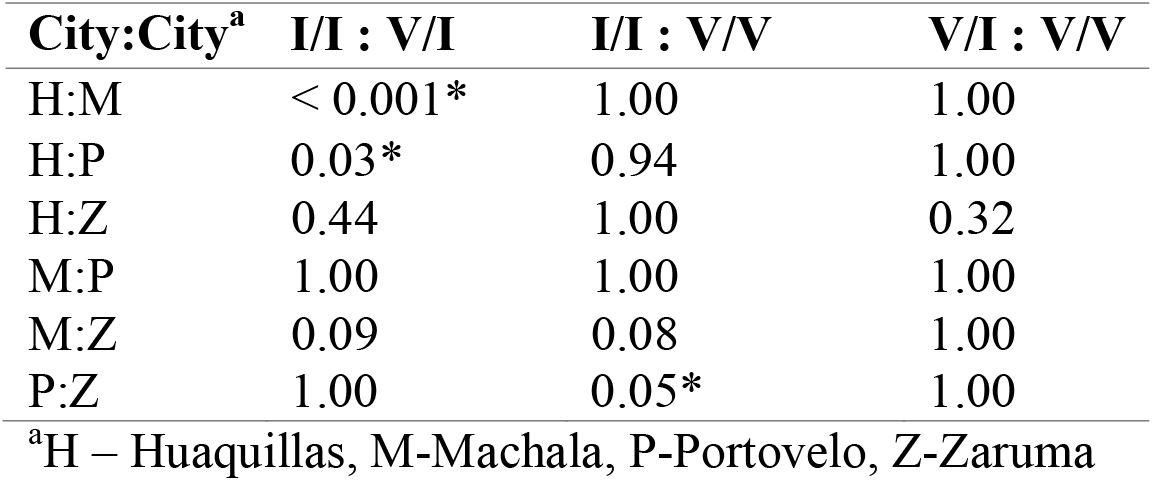
Post-hoc Fisher’s exact test *p*-values for genotype V1016I in season 1, with comparisons in genotype frequencies made between the cities of Huaquillas (H), Machala (M), Portovelo (P), and Zaruma (Z). Significant difference are denoted with an asterisk.

| **City:City<sup>a</sup>** | **I/I : V/I** | **I/I : V/V** | **V/I : V/V** |
| --- | --- | --- | --- |
| H:M | < 0.001* | 1.00 | 1.00 |
| H:P | 0.03* | 0.94 | 1.00 |
| H:Z | 0.44 | 1.00 | 0.32 |
| M:P | 1.00 | 1.00 | 1.00 |
| M:Z | 0.09 | 0.08 | 1.00 |
| P:Z | 1.00 | 0.05* | 1.00 |

<sup>a</sup>H – Huaquillas, M-Machala, P-Portovelo, Z-Zaruma

**S7 Table:** Post-hoc Fisher’s exact test *p*-values for genotype F1534C in season 1, with comparisons in genotype frequencies made between the cities of Huaquillas (H), Machala (M), Portovelo (P), and Zaruma (Z). Significant difference are denoted with an asterisk.

| **City:City<sup>a</sup>** | **C/C : F/C** | **C/C : F/F** | **F/C : F/F** |
| --- | --- | --- | --- |
| H:M | 1.00 | 1.00 | 1.00 |
| H:P | 1.00 | 1.00 | 1.00 |
| H:Z | 0.02* | 1.00 | 1.00 |
| M:P | 1.00 | 1.00 | 1.00 |
| M:Z | < 0.001* | 1.00 | 1.00 |
| P:Z | 0.18 | 1.00 | 1.00 |

<sup>a</sup>H – Huaquillas, M-Machala, P-Portovelo, Z-Zaruma

**S8 Table:** Post-hoc Fisher’s exact test *p*-values for genotype V1016I in Machala, with comparisons in genotype frequencies made between seasons. Significant difference are denoted with an asterisk.

| **Season:** | **I/I :** | **I/I :** | **V/I :** |
| --- | --- | --- | --- |
| **Season** | **V/I** | **V/V** | **V/V** |
| 1:2 | 1.00 | 1.00 | 1.00 |
| 1:3 | 1.00 | 0.02 | 1.00 |
| 2:3 | 0.02* | 0.43 | 1.00 |

**S9 Table.** Post-hoc Fisher’s exact test *p*-values for genotype V1016I in Portovelo, with comparisons in genotype frequencies made between seasons. Significant difference are denoted with an asterisk.

| **Season:**<br>**Season** | **I/I :**<br>**V/I** | **I/I :**<br>**V/V** | **V/I :**<br>**V/V** |
| --- | --- | --- | --- |
| 1:2 | 0.05* | 0.36 | 1.00 |
| 1:3 | 0.001* | 0.005* | 1.00 |
| 2:3 | 1.00 | 0.58 | 1.00 |

**S10 Table.** V1016I and F1534C Genotypes compared to deltamethrin-resistant phenotype

| V1016I | I/I<br>(mutant) | V/I<br>(heterozygote) | V/V<br>(wild type) | *p*-value<sup>a</sup> |
| --- | --- | --- | --- | --- |
| *Resistant* | 107 | 79 | 10 | <0.001* |
| *Susceptible* | 17 | 88 | 52 |  |
| F1534C | C/C<br>(mutant) | F/C<br>(heterozygote) | F/F<br>(wild type) |  |
| *Resistant* | 190 | 5 | 1 | 0.10 |
| *Susceptible* | 144 | 10 | 3 |  |

<sup>a</sup>Fisher's Exact Test

**S11 Table.** V1016I and F1534C Genotypes compared to alpha-cypermethrin resistant phenotype

| V1016I | I/I<br>(mutant) | V/I<br>(heterozygote) | V/V<br>(wild type) | *p*-value<sup>a</sup> |
| --- | --- | --- | --- | --- |
| *Resistant* | 80 | 58 | 2 | <0.001* |
| *Susceptible* | 23 | 84 | 32 |  |
| F1534C | C/C<br>(mutant) | F/C<br>(heterozygote) | F/F<br>(wild type) |  |
| *Resistant* | 138 | 2 | 0 | 0.10 |
| *Susceptible* | 132 | 7 | 0 |  |

<sup>a</sup>Fisher's Exact Test

**S12 Table.** Exact test *p*-values for the exact test of Hardy-Weinberg equilibrium for the V1016I genotype frequencies in each city and season. Significant p-values indicating the population is likely not in Hardy Weinberg equilibrium are denoted with an asterisk.

| **City** | **Season** |  |  |
| --- | --- | --- | --- |
|  | **1** | **2** | **3** |
| Huaquillas | 0.03* | 0.10 | 0.73 |
| Portovelo | 1.00 | 0.06 | 0.02* |
| Machala | 0.03* | 0.63 | 0.18 |
| Zaruma | 0.11 | na | na |

**S1 Figure:**
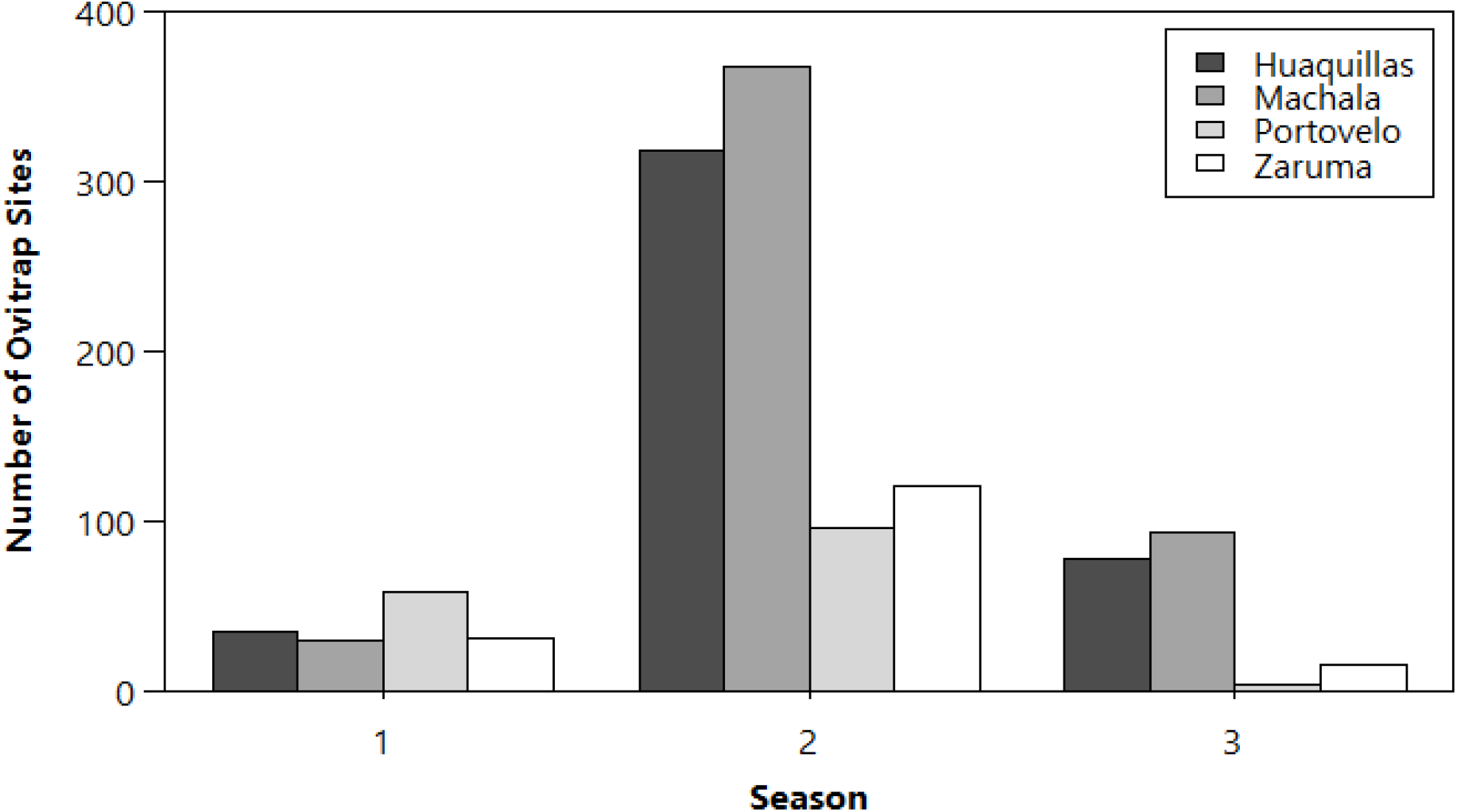
Dengue cases by epidemiologic week for all study cities during seasons 1-3. Total weekly dengue cases reported to the Ecuador Ministry of Health are given for Huaquillas, Machala, Portovelo, and Zaruma

**S2 Figure:**
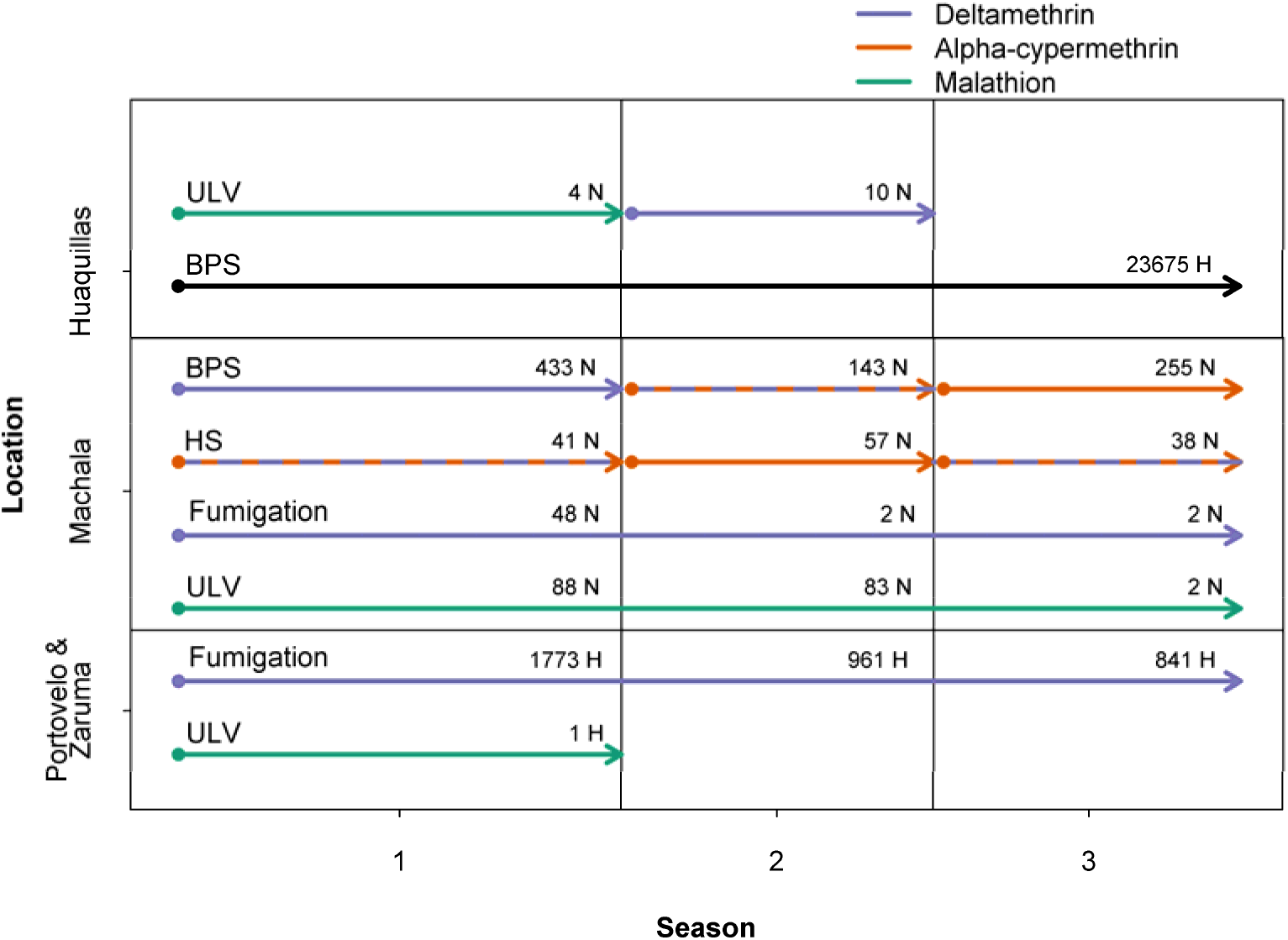
Ovitraps by city and season.

**S3 Figure:**
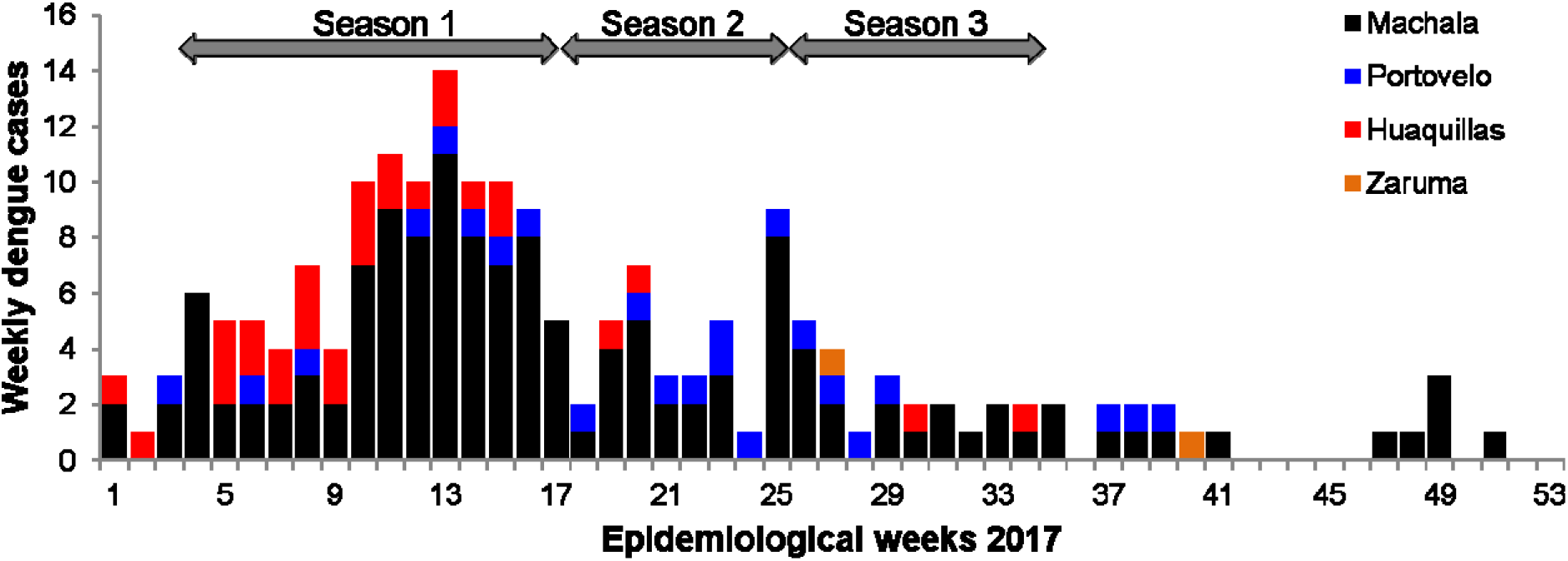
Genotypic characterization of the V1016I mutation. Dissociation curves representing the susceptible (wild type) homozygous (A), heterozygous (B), and resistant (mutant) heterozygous (C) genotypes.

**S4 Figure:**
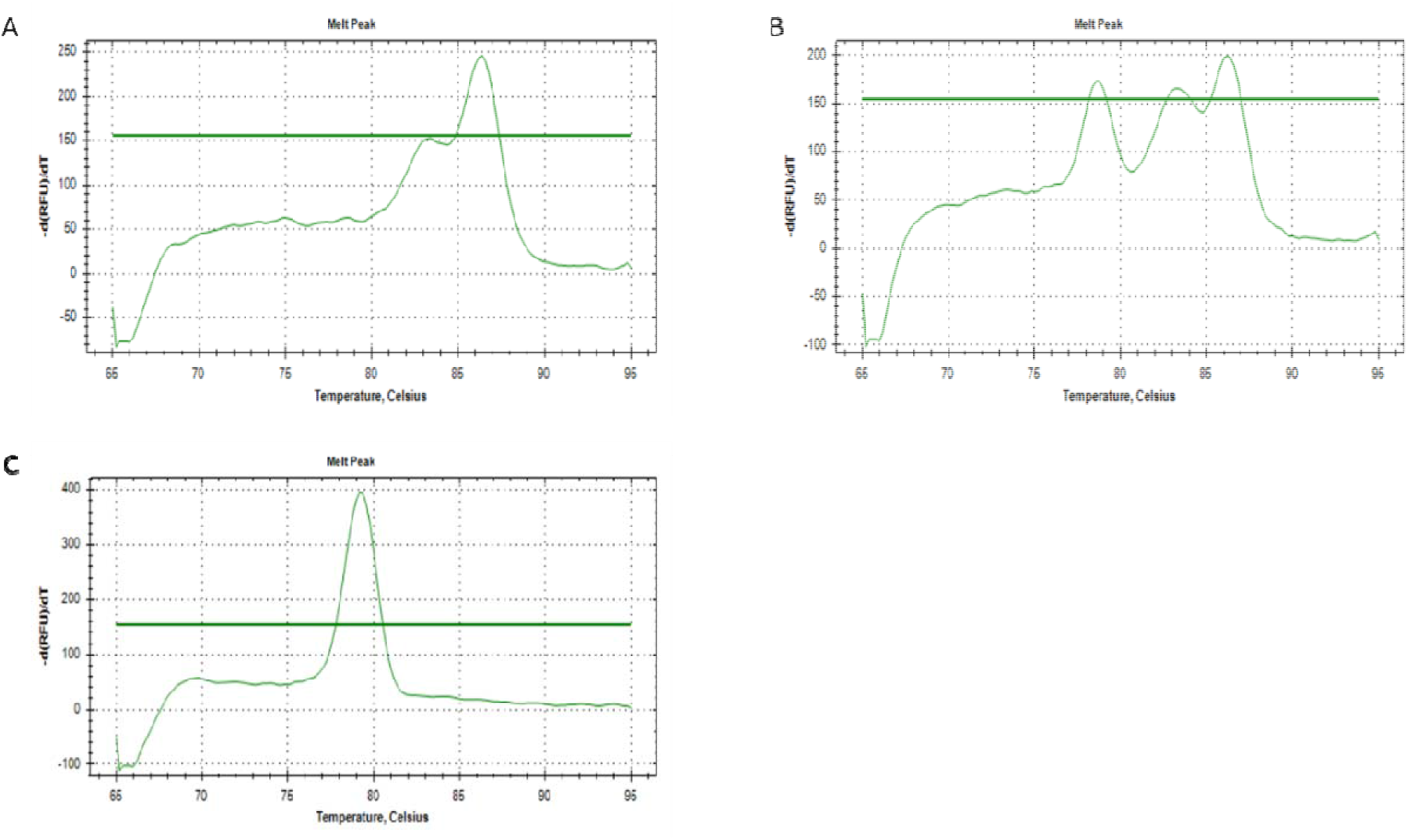
Genotypic characterization of the F1534C mutation. Dissociation curve representing the wild type (susceptible) homozygous (A), heterozygous (B), and mutant (C) genotypes.

**S5 Figure:**
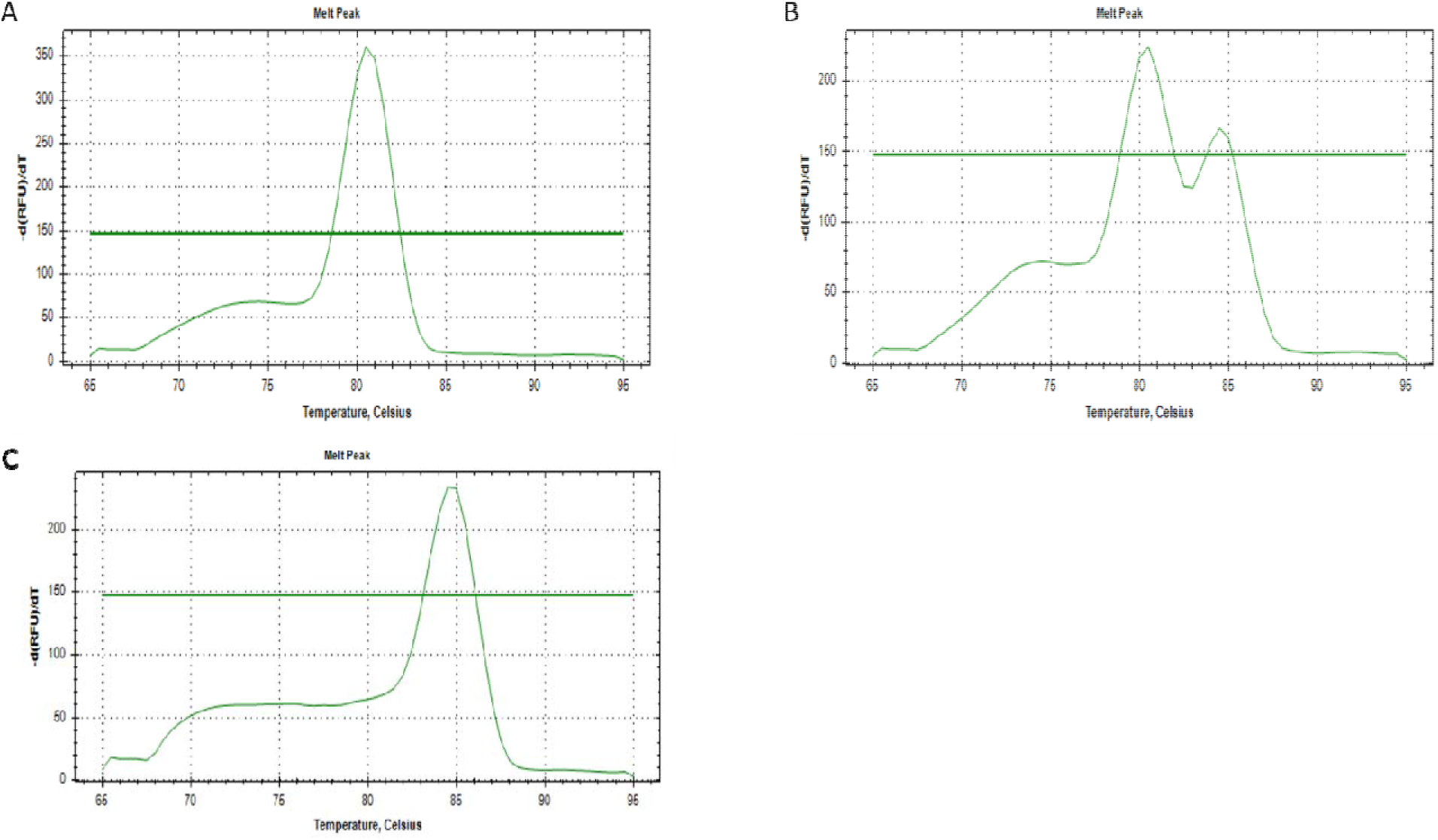
City-wide insecticide application. Timing, application method, and number of houses or neighborhoods treated are shown. Dashed arrows represent a mixture of two insecticides. One insecticide product (black) was unidentified. The house number for Huaquillas represents total houses treated over the season. ULV=ultra low-volume spraying, BPS=backpack spraying, HS=handpump spraying, H=houses, N=neighborhoods.

